# Maximum Likelihood Estimators For Colony Forming Units

**DOI:** 10.1101/2023.05.18.541301

**Authors:** K. Michael Martini, Satya Spandana Boddu, Ilya Nemenman, Nic Vega

**Author notes:** **For correspondence:**, (KMM); (NV).

## Abstract

There is a recognized need to measure the abundance of microbes in hospital environments, in the sanitation industry, and in food preparation. Doctors, microbiologists, and food safety experts have been addressing this need by using serial dilution methods to grow bacterial colonies in small enough numbers to count and, from these counts, to infer bacterial concentrations measured in Colony Forming Units (CFUs). There are two primary types of such methods: plating bacteria on a growth medium and counting their resulting colonies or counting the number of tubes at a given dilution that have growth. Traditionally, these types of data have been analyzed separately using different analytic methods. Here we build a direct correspondence between these approaches, which allows one to extend the use of the Most Probable Number (MPN) method from the liquid tubes experiments, for which it was developed, to the growth plates. We also discuss how to combine measurements taken at different dilutions, and we review several ways of analyzing colony counts, including the Poison and truncated Poison methods. For all methods, we discuss their relevant error bounds, assumptions, strengths, and weaknesses. We provide an online calculator for these estimators.

## Introduction

Extrapolation of viable microbial counts from suspensions of live cells is a longstanding—and surprisingly complicated—problem. The fundamental problem is simple: there exists a volume *V*_0_ with some unknown concentration of live microorganisms, which an experimentalist wants to measure. That initial volume will be serially diluted (usually in a ten-fold series), and fixed-volume aliquots (sub-samples) of the resulting suspensions will be cultured. If these aliquots are spread or dropped onto agar plates, the resulting data will be in the form of colony counts. Alternately, multiple aliquots may be taken from a single dilution and used to seed a number of wells or tubes of liquid culture, or a number of plates. Then the number of volumes showing growth when seeded from a particular dilution, as a fraction of the total number of volumes inoculated, can be used to calculate the Most Probable Number (MPN) of live agents in the initial volume (***McCrady, 1915; Cochran, 1950***).

Best practice for this apparently simple and ubiquitous scenario has been the subject of debate for over a century (***McCrady, 1915; Breed and Dotterrer, 1916; Fisher et al., 1922; Halvorson and Ziegler, 1933; Jennison and Wadsworth, 1940; Jones et al., 1948; Skinner et al., 1952; Johnson and Brown Jr, 1961; Taylor, 1962; Harris and Sommers, 1968***). There are technical considerations to this problem. For example, all plates or tubes used for growth should have the same ability to support growth of the organism(s) being studied, and the sample must be sufficiently homogenized to ensure that microbes are free in solution and not adhered to one another or to a substrate (***Fisher et al*., *1922***). However, such considerations are case-specific and beyond the scope of the present work.

These counts are subject to counting errors as well. At one extreme, when the sample is too concentrated, the number of resulting colonies will be too numerous to count (TNTC; sometimes “too many to count”, TMTC). At these high concentrations, colonies merge, breaking the assumption that each microbe corresponds to one colony (***Jennison and Wadsworth, 1940; Ben-David and Davidson, 2014***). At the other extreme, when the sample is very diluted, the number of colony initiating bacteria in the sample is subject to small-number statistical (sampling) fluctuations, resulting in high relative error (ratio of the standard deviation to the mean) (***Haas et al., 2014; Christen and Parker, 2020***). Finally, experimental errors, such as inaccuracies in pipetting, can emerge and compound over the steps of a serial dilution. However, the latter source of error is expected to be negligible for equipment calibrated to usual standards, and technical replication further reduces effects of this variation (***Hedges, 2002***).

Thus the problem at hand is: How can CFU density best be estimated from plate counts, given the error produced by sampling fluctuations, colony crowding, and (to a lesser degree) pipetting? These errors will contribute differently to different experimental designs. For a *single sample represented by one count of colonies n*_*k*_ at one dilution *d*_*k*_ (because only one dilution was measured, or because only one spot or one plate in a series was countable), statistical error of counts (presumably Poisson) is inevitable, and pipetting error will contribute but may not be significant. For a *single sample represented by more than one count of colonies* (representing counts at different dilutions within a single dilution series, and/or technical replicates where one sample was measured multiple times), the same errors apply, but pipetting bias may not be constant across measurements (for example, one failing O-ring on a multichannel pipette can lead to bias in a single column of a 96-well plate).

For *multiple samples of the same type measured in parallel* (biological replicates), we can no longer expect variation across samples to reflect a Poisson-distributed sampling error. Indeed, individual measurements will be subject to sampling variation, but variation across samples will be biological (or otherwise inherent), and demographic (accumulating over time) in addition to sampling. This was the basis for the Nobel-prize winning experiments of Luria and Delbruck, who used the distribution of fluctuations to distinguish Darwininan vs. Lamarckian evolution (***Luria and Delbrück, 1943***). This is also frequently the case in environmental samples, where different samples from the same source (e. g., water samples from different parts of the same lake) will produce measurements that have super-Poisson variation (aka, over-dispersed). In such cases, some of the variation is “real” due to inhomogeneities in the source, and it cannot be modeled as mere sampling error. Biological variation is problem dependent and often carriers in it the imprint of the underlying fundamental biology (***Luria and Delbrück, 1943***); it will not be dealt with here. Instead, we will focus on estimation of CFU density within an individual sample, which may be represented by a single set of measurements or by technical replicates, in which one sample is measured multiple times.

The main objective of this paper is to propose methods for accurately estimating colony forming units (CFUs), while taking into account the effects of crowding and sampling fluctuations, without losing valuable data from counts. Drawing from previous research (***Haas et al., 2014; Ben-David and Davidson, 2014; Sutton, 2011; Christen and Parker, 2020***), we present simple analytical formulas that can be used to combine counts from different dilutions and to obtain precise CFU estimates along with accurate error bars. First, we examine existing methods in the literature, assessing their strengths and weaknesses. Next, we introduce the “Poisson with a cutoff” method, which clarifies the impact of crowding on CFU density estimation and demonstrates how to minimize the effects of sampling error by combining measurements of “uncrowded” counts. Finally, we use a crowding-explicit model to demonstrate the relationship between canonical plate-based counts and the Most Probable Number method for presence/absence of growth in liquid media. We conclude by providing practical recommendations for experimentalists on how to select appropriate dilution and replication schemes and how to combine data from multiple observations. We also have provided a calculator for these estimators available on `Hugging Face` spaces, named `CFUestimator` (***Martini, 2023***).

## Results

### A Brief History of Counts

Colony Forming Units (CFUs) are a proxy for the concentration of microbes within a sample. The experimental procedure for estimating CFUs consists of serially diluting homogeneous samples in a sterile aqueous buffer, then plating aliquots of these dilutions on growth-supporting agar and later counting the resulting colonies. If an appropriate dilution has been reached, each microbe will form an independent colony that is countable by eye. The simplest way of estimating CFUs is to multiply the number of colonies by the reciprocal of the dilution factor to find the concentration of colony-forming microbes in the original suspension (***McCrady, 1915; Halvorson and Ziegler, 1933; Sutton, 2011***). For example, say there is a single sample represented by one countable 10 cm plate in a dilution series, where we observe 100 distinct colonies after plating 100 *μ*L of a 1:100 dilution (dilution 2 in a ten-fold series) from the original sample. In this case, following this simple procedure, we would obtain:

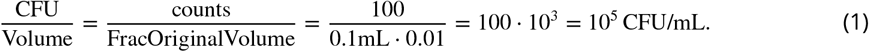

This is exactly equivalent to multiplying the number of counts by a volume correction factor (1/(size of aliquot in mL)) and multiplying by the base of the dilution series raised to the power of the number of dilution steps:

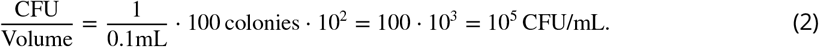

This simple calculation follows from a more general Poisson model, explained below. This method works reasonably well under ideal conditions: all samples should be represented by a single count of colonies, and each count should be large enough to minimize small-number sampling fluctuations, and yet small enough to avoid crowding on the plate. When any of these conditions are not met, accurate estimation of CFU density becomes more complicated.

There is a broad literature of methods proposing to ensure that estimates of CFU density are “good”. A *good* estimator should be accurate. Formally, this means that such estimators should have the true value of the CFU density as their expected value. In other words, they must be unbiased. *Good* estimators must also be precise, so that variance in the estimate is small and samples are repeatable. Therefore, an ideal solution to this problem should provide an estimator that is provably unbiased and with a minimal variance. The solution to this problem is well known in statistics: if we can assume that data follows a specific probability distribution, then the *maximum likelihood estimator (MLE)* for that distribution will have these properties (***Haas et al., 2014***). While this is formally true only for very large samples, MLE estimators generally perform well even for small samples. Further, an ideal method should be straightforward to use in the hands of researchers without advanced mathematics background. Unfortunately, many of the available methods fail one or the other of these requirements, being either simple to use, but statistically sub-optimal, or mathematically correct, but incomprehensible to many bench scientists.

Straight-forward to use methods focus largely on designing protocols that avoid data in error-prone extremes. For example, the FDA recommends (***Blodgett, 2020***) that the best dilution range for coliform bacteria results in 25 to 250 colonies per 10-cm plate, with the ideal count closest to 250. Restriction on the high end limits counting errors due to growth limitation by nutrient depletion as well as outright merging of colonies, which would bias the number of counts downward. Conversely, restriction on the lower end limits the sampling error associated with small numbers of counts. Specifically, under the assumption that counts represent random draws from a given sample and are, therefore, Poisson-distributed, the error scales as the square root of the number of counts. Thus, for small counts, the error becomes an unacceptably large fraction of the mean. Within the example above, our dilution 2 count of 100 colonies should have a standard deviation (SD) of 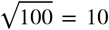, giving a coefficient of variation (CV) of 10%. At dilution 3, we might obtain 10 counts, with a SD of 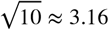, and a CV of 31.6%.

From here, the simplest approach that is often used in practice is to choose only the plate or spot that has the “best” count in the acceptable range, and to estimate CFU density based on that single count. Often only the dilution at the high end of the countable range is used since it has the smallest sampling fluctuations; all other measurements are discarded (***Sutton, 2011***). We call this the “pick-the-best” method for later reference. If counts in the acceptable range can be consistently achieved, this method is straightforward and reasonably accurate. However, discarding data is rarely advisable, and over- and under-crowded measurements can, in fact, be used to improve CFU estimates.

### Simplest “Good” Estimator: Poisson

One simple and reasonably accurate model for calculating CFUs assumes that the number of colonies are Poisson distributed, with variation due to sampling. That is, for a particular dilution, the mean colony count for that dilution is the same as the variance. This model ignores crowding effects but works well for modeling sampling fluctuations. By this model, the most likely estimator for the density of microbes is simply the ratio of the total number of colonies counted from all plates divided by the total amount of liquid used from the original sample in all plates (see *Supplementary Information*). If there is only one countable measurement for a given sample, this simplifies to “pick-the-best”.

The Poisson model implicitly assumes that the original sample is well mixed and each microbe plated will result in its own separate and countable colony. It further assumes that experimental volume is spread uniformly across the agar surface, resulting in cells being randomly distributed, independent of the locations of where other cells landed. Formally, these assumptions mean that there is a uniform and well mixed population density *r* of microbes per unit volume in an initial volume of liquid *V*. The liquid is diluted by a factor *d*_*k*_ = *V*_*k*_/*V*, where *V*_*k*_ is the volume of the liquid from the original sample used on the plate or the spot *k*. Plating will result in *n*_*k*_ colonies, where *n*_*k*_ is Poisson distributed with the parameter *λ* = *rd*_*k*_*V* = *rV*_*k*_. That is, the average number of colonies per experiment is *rd*_*k*_*V* with variance *rd*_*k*_*V*. Using these assumptions, the MLE estimator of the density of microbes *r*_mle_ and its standard error are:

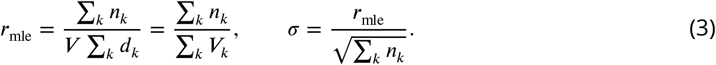

In other words, the best estimator for the concentration, *r*_mle_, is the total number of colonies divided by the total amount of the original volume of liquid used. However, as noted earlier, this ignores crowding and counting errors. In practice, this method should be avoided unless all measurements are from well-dispersed, uncrowded plates, as crowding effects can make a large difference in the estimator, resulting in underestimating the microbial density as colonies merge and are under-counted.

If technical replicates exist (multiple measurements of the same sample), it is straightforward to test whether the data adhere to a Poisson distribution using the following test, known as the dispersion index test. If there are *j* measurements of a given sample, with average number of counts 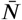 and variance of counts 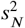, then the index of dispersion *D*^2^ is:

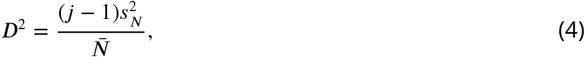

which will be distributed as *χ*^2^ with *j* − 1 degrees of freedom (***Haas et al., 2014***). If *D*^2^ is greater than the upper 1 − *α* quantile of that distribution, where *α* is the needed significance p-value, then we reject the null hypothesis that these replicates are drawn from the same Poisson distribution. This can indicate technical problems that are introducing an excess of variation, possibly by biasing replicates differently from one another (e. g., the failing O-ring example above), or biases due to a too-lenient cutoff for countability.

### Combining Data: Common Bad Estimators

The primary reason for the “pick-the-best” approach is that it eliminates confusion over how to combine multiple measurements for a given sample, particularly when counts belong to more than one dilution. First notice that combining measurements from technical replicates that are taken at the same dilution is straightforward. For example, let’s assume an original 200 *μ*L volume *V* contains *r* = 3 · 10^8^ CFU. We can create simulated serial dilutions from this original volume by assuming that each pipetting step (ten-fold dilutions and plating onto agar) is a binomial sampling event (***Christen and Parker, 2020***) that comes with experimental noise. In one such simulation, triplicate plating 100 *μ*L aliquots results in counts *n*_6_ =(162, 141, 148), all from the sixth ten-fold dilution. The fraction of the original volume plated in each case is *V*_6_ = 0.5 · 10^−6^ = 5 · 10^−7^. These numbers can be combined via the Poisson method shown in the previous section to estimate CFU density in *V* :

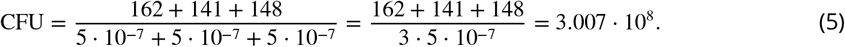

Alternately, counts taken from the same dilution can be averaged across technical replicates, then adjusted by the volume plated and the dilution read (***USDA, 2015***):

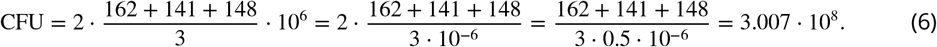

Clearly, these two most common approaches are algebraically identical.

In contrast, combining counts across different dilutions is less straightforward. In fact, some commonly-used methods for combining measurements are statistically inadmissible. For example, if there are multiple measurements in the countable range, the USDA recommends (***USDA, 2015***) that researchers calculate the estimated CFU for each dilution separately using the average colony count across technical replicates at a given dilution and then average the results of the separate dilutions. If the two estimates are more than a factor of 2 apart, the researcher is instructed to instead only use the counts from the higher-density plates. This commonly used method, incorrectly combines the data using a simple average, thus increasing the variance of the estimated CFU density. Indeed, continuing the example above, let’s suppose that, on the plates corresponding to the seventh ten-fold dilution from these three technical replicates, we observe (13, 17, 20) colonies. The Poisson estimator gives us:

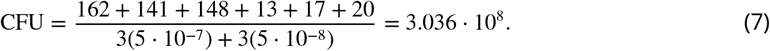

The USDA averaging method gives:

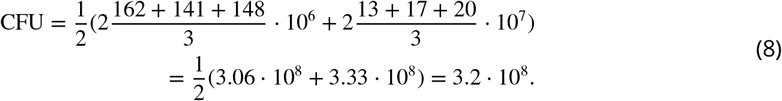

On these data, averaging was substantially less precise, with an error of 7% as compared with the Poisson method’s error of 1% (recall that the true density in this simulated example is 3.0 · 10^8^ CFU per 200 *μ*L). The USDA method improperly averages across dilutions, amplifying fluctuations associated with small colony number counts, whereas the simple Poisson model properly combines measurements across dilutions by effectively re-weighting small counts by the small volumes with which they are associated. In a later section, we demonstrate that averaging across dilutions will, as a rule, increase the variance of CFU estimates.

### Too Few and Too Many

Further, there is the problem of what to do with zero counts. These data are inevitably limited by some threshold of detection (TOD), representing the smallest CFU density at which counts can be detected. This “left-censoring” is a well-known issue (***Canales et al*., *2018; Gijbels, 2010; EPA, 2014***) with many proposed work-arounds, including but not limited to: substituting zeros with a small value (which may be the average of the undetectable range, a maximum-likelihood inferred value, or some other small number), reporting zeros as “below TOD” or “<1” rather than as a value, and pretending they didn’t happen (not generally recommended; although if zeros are rare, it won’t make much difference) (***Canales et al*., *2018; EPA, 2014***). Sometimes, a threshold of quantification (TOQ) representing the lowest “acceptable” (sufficiently precise) count is used along with or instead of TOD (***Sutton, 2011***), with values below this threshold omitted from analysis.

The “best” approach to zero-contaminated count data depends on what else is in the data and what the data will be used to do. If a sample is represented by zero and non-zero measurements, the Poisson model explicitly allows zero counts to be incorporated as outcomes of the random sampling process. For example, if a hypothetical *V* = 200 *μ*L sample contains 5 · 10^7^ CFU, one simulation of serial dilution and plating in triplicate with 100 *μ*L per plate produces dilution-6 counts of (31, 26, 20) and dilution-7 counts of (4, 0, 0). Using just the dilution-6 counts, we estimate

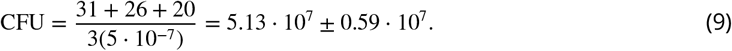

If we use the lower dilution as well, we obtain

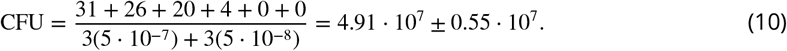

In this case, incorporating data from zeros (in the form of the additional volume that was plated but contained no counts) improved precision. Alternately, when zeros are common because the density in the original sample is close to the TOD, non-zero counts are useful for making a distinction between samples where no organisms are detectable (and density might be zero) and those where the density of organisms cannot be zero. Although the actual density cannot be estimated accurately or precisely from very low counts, the distinction between “<TOD” and “>1” for a given sample is important (***EPA, 2014***).

At the other end of the range, researchers must deal with crowding and set thresholds for “too many to count”. Defining an optimal range for “countable” data is not always straightforward, and this determination is very important to ensure that CFU estimates are accurate. Since the sampling-based standard error of counts scales as 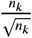, the number of colonies counted *n*_*k*_ should be as large as reasonably possible.

However, there are consequences for pushing this too far. As cell density in the aliquot increases, counts will be biased downwards due to merging of colonies and colony stunting or failure to grow. These data are then “right-censored”, with an upper limit past which the number of counts observed does not increase in proportion to an increase in the density in the original sample. Densities above this point result in “crowded” samples, with counts that are lower than the true number of colony forming units. Further, as the number of colonies per plate or spot increases, data collection becomes more time-consuming; it is common for researchers to minimize effort on plates near the top of the “acceptable” range by dividing plates into sections, counting colonies in one section, and multiplying this count by the number of sections to get an estimated final count for the whole plate (***Blodgett, 2020***). While this approach is sufficient for a rough estimate of CFU density, it introduces additional sampling variation due to both reduction in counts and imperfect division of plates, and it does not remove bias due to crowding. We will demonstrate the consequences later in this paper.

Previous works (***Ben-David and Davidson, 2014***) have modeled crowding using shifted Poisson distributions. In these models, the variance of estimates from crowded data would be the same as if there was no crowding and the mean would be shifted down due to colonies merging together. However, this is *a priori* unlikely to be true. As we will show below, if colonies are crowded, both the mean and the variance will be shifted relative to the pure Poisson (uncrowded) distribution. The reason for this is that the variance of the large colony counts is shifted downward due to a “ceiling” effect—there is an upper bound to the total number of colonies, which limits upwards fluctuations. In other words, the use of a shifted Poisson distribution is a reasonable approximation, but the variance must also be modified.

### Better Estimators: Poisson With Cutoff, aka What’s Countable, Exactly?

The main problem with the naive Poisson model is that it does not account for counting errors due to crowding. The simplest way to take account of the crowding is to assume that there is a threshold of colonies, *M*, below which crowding is negligible, which in practice will often be smaller than the largest number of counts we are willing to attempt. We can then segment our data into two parts: plates with counts above the threshold where crowding is important, and plates with counts below the threshold where crowding is not important. If we have identified our cutoffs well, the naive Poisson estimator above is correct for all measurements where the number of colonies counted *n*_*k*_ ≤ *M*. The calculation is, therefore, exactly the same as for the naive Poisson estimator given above, with the caveat that only measurements *n*_*k*_ ≤ *M* are used. Here, the indicator function *I*(*n*_*k*_ < *M*) is 1 when *n*_*k*_ < *M*, and 0 otherwise. Similarly, *I*(*n*_*k*_ > *M*) is 1 when *n*_*k*_ > *M*, and 0 otherwise.

Due to its balance between simplicity and accuracy, this method is the easiest to use in practice.

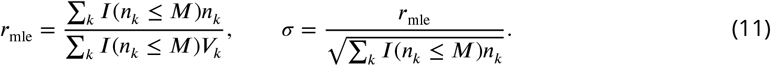

If we want to incorporate data from measurements above this threshold *M*, the calculation becomes slightly more complicated. Using “crowded” measurements as if they were uncrowded will bias the naive Poisson estimator downward, resulting in under-estimation of CFU density (Fig. 2). In a *sophisticated* version of this model, we can use the number of plates/spots that were above the crowding threshold *M*, along with the colony counts from plates/spots below this threshold at the same dilution, to estimate CFUs. This will be applicable when plate counts at a given dilution are toward the high end of the countable range, such that some technical replicates fall below this threshold and others above it by chance. To estimate the CFU density in the original sample *r*, the following equation should be solved numerically (see SI for the derivation):

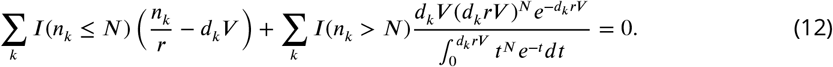

The first term is equivalent to the simple Poisson model and uses the counts from uncrowded samples directly, whereas the second term reflects the probability of counts being above the threshold *M*. Inference of *r* can be done in Excel using SOLVER or using numerical solvers in R, Python, MATLAB, etc. An equivalent model is shown in (***Haas et al., 2014***).

This model properly accounts for two error sources: the sampling fluctuations and the crowding effect. The simple Poisson, using only counts from uncrowded plates, gives a good estimate for the CFU counts and properly combines multiple measurements at different dilution factors. The more sophisticated form of the model has greater precision, but the greater computational effort may or may not be worth it to an investigator depending on the effect size and the structure of the experiment. In the next section, we present an alternate estimator based on the Most Probable Number approach, which we argue provides a better trade-off between effort and estimator performance when incorporating data from crowded samples.

### Crowding and the Most Probable Number

For the final model we consider the effects of crowding in space. To account for crowding, we will divide each plate into 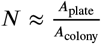 regions, each approximately the size of a full colony. We make the assumption that if more than one microbe lands in one of these regions, the colonies that form from these cells will grow together and be counted as one colony. For each region, the number of cells landing in that region will be Poisson distributed with parameter 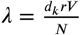.

These assumptions are equivalent to that of quantal-based methods for microbial quantification, such as the commonly used Most Probable Number (MPN) method. In the MPN assay, a known quantity (volume of original sample) is introduced into each of a series of *N* replicate tubes, and the dilution of the original sample is adjusted to find a region where some of the tubes contain viable growth and some do not. The results of this assay are therefore, for each dilution volume *V*_*k*_ from the original sample, out of the *N*_*k*_ tubes inoculated, a number *n*_*k*_ that is positive for growth.

A direct mapping to tube-based assays is possible if space on a plate (or within a spot) is considered as a set of colony-sized bins. Each of the N colony-sized regions on a plate or within a spot corresponds to one tube. The presence of colonies in a particular region corresponds to when a tube has growth. Hence a plate that is divided into *N* regions can be thought of as *N* tubes being tested in parallel, cf. Figure 1.

**Figure 1.**
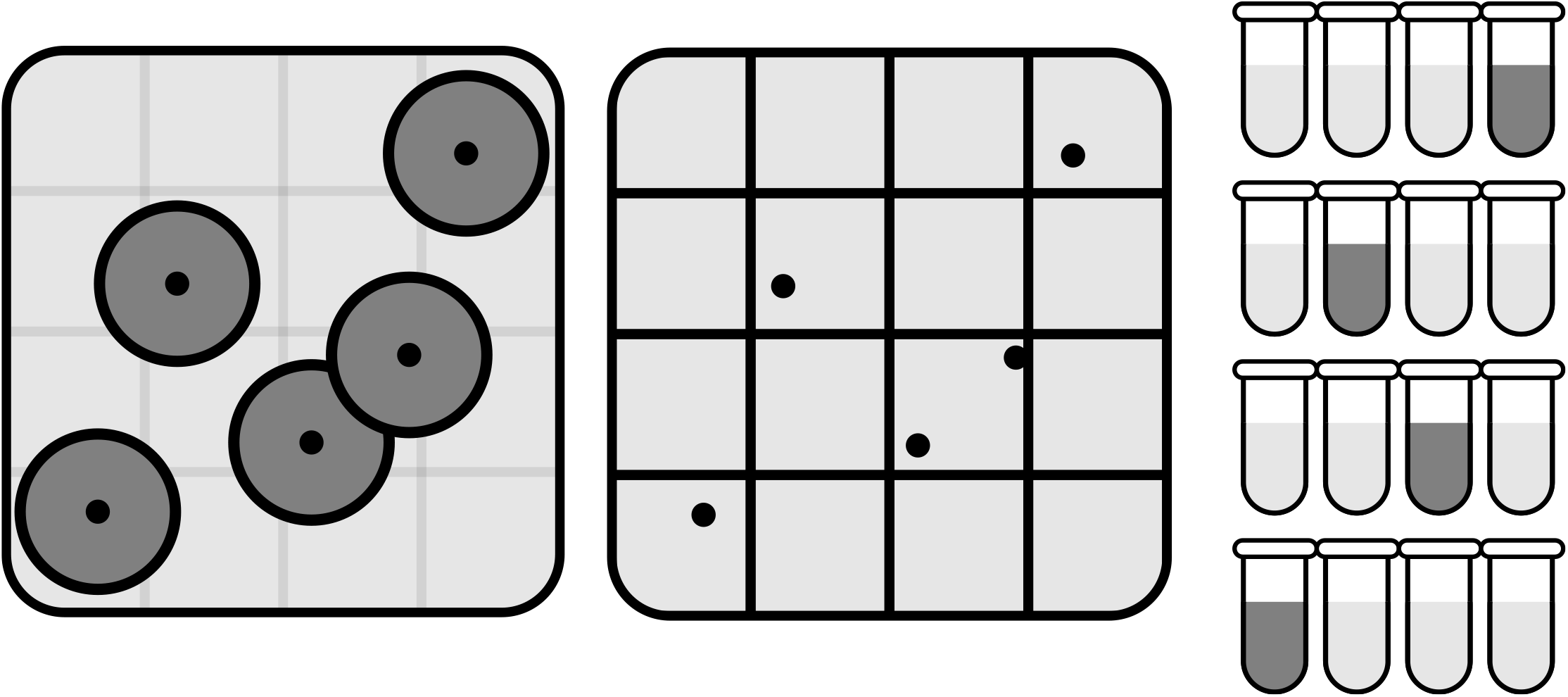
Visual equivalence between plate and tube based assays. The left panel is a cartoon of a typical plate containing colonies, where the growing colonies are shown as dark disks. In the middle panel, the plate is divided into *N* (here 16) approximately colony-sized regions. If a region contains one or more colony centers (black dots), this region can be mapped to a positive (dark) tube as shown in the right panel. Similarly regions containing no colony centers are mapped to negative (light) tubes. This demonstrates that plating is equivalent to a massive parallel version of a tube based assay with 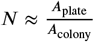 tubes. Furthermore it demonstrates that the MPN method can be used for plate data.

**Figure 2.**
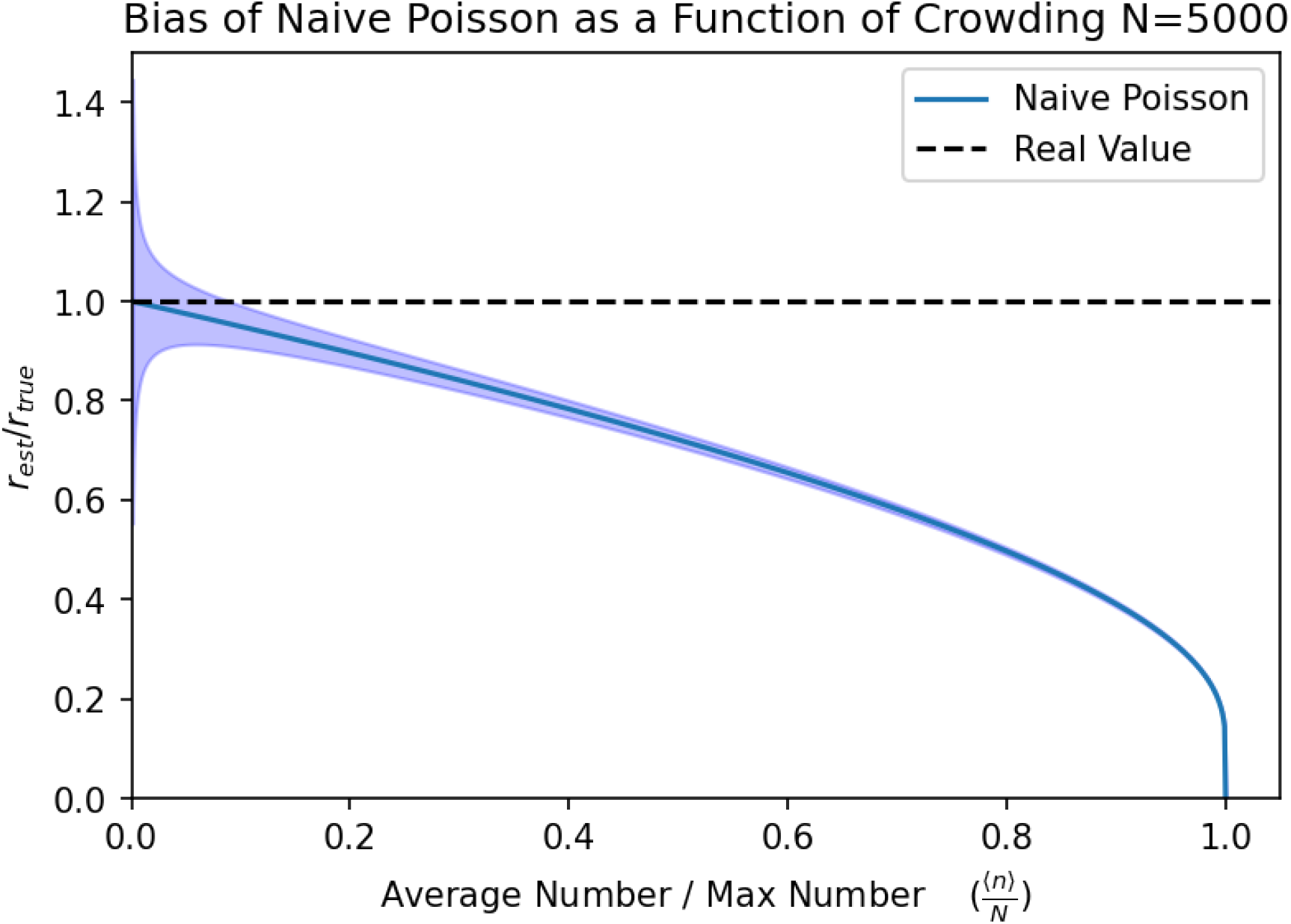
The naive Poisson estimator underestimates the true concentration and becomes more biased as a function of crowding. We illustrate this by plotting the ratio of the estimated concentration (with the error bands denoting ± one s. e. m. at *N* = 5000) to the true concentration. Here crowding is measured by the ratio of the average number of colonies to the maximum number of colonies that can fit within a plate 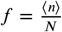. At low crowding values, the naive estimator has low bias, but large uncertainty. At a crowding value of 0.2 the naive-Poisson estimator underestimates the true concentration by about 10%, and many-fold underestimation is possible as crowding approaches 1.

Therefore, the probability of *n*_*k*_ successes in *N* colony-size regions on the agar surface can be described using a crowding-explicit model based on the binomial distribution. Assuming that the cells in the original sample are well-mixed, the probability of zero cells landing in a particular region is (from the Poisson) 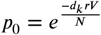 and the probability that at least one cell lands in that region is therefore 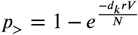. Assuming that the original sample is well mixed, each region is independent of all other regions in our crowding model, so that

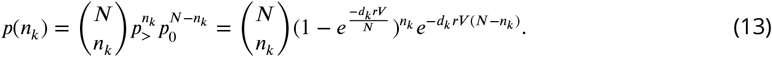

We can maximize this probability to find the MLE CFU density, *r*_mle_ (see the SI fo the full derivation). We can accomplish this by numerically solving the following equation for *r*:

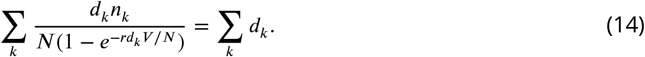

This expression for *r* is the same as that of the MPN estimator(***Blodgett, 2020; Alexander, 1983***). In the Appendix we show that, in the limit where concentrations and colony counts are low, this model simplifies to the Poisson model. Outside the “uncrowded” regime, the mean and the variance of data from the crowding model are not the same as in the Poisson. Therefore, the two approaches are not equal to each other, though both are depressed due to the “ceiling” effect described earlier. In the Appendix, we also find that the error associated with the maximum likelihood estimator *r*_mle_ of the MPN method can be minimized at an optimal dilution factor, which falls into the crowded regime.

The MPN procedure can generate biased estimates of the original sample density, and the precision and accuracy of results depend strongly on the number of tubes used (***Haas et al., 2014***). The bias on the maximum likelihood estimator results in an over-estimate of 20-25% with 5 tubes, which is reduced to a few percent with 50 tubes (see Appendix). By back of the envelope calculation, an average 10 cm plate (inside diameter 86 mm, surface area 58 cm^2^) can fit a maximum of approximately 5000 medium-sized (1 mm outside diameter) “tubes”, whereas a single grid square on a 10 × 10 cm square plate (typically gridded 6 × 6) can fit 200 of these colony-sized spaces. All of these are well above the threshold where the bias in this estimator (***Salama et al*., *1978***) makes much difference in the value. (Note that this refers to the number of *colony-sized spaces* available and is independent of the number of colonies observed.) This also means that the standard error of the estimator will, in theory, be minimized at a plating density that is much higher than the threshold for “uncrowded” plates and, in fact, is well into a range of densities where a minority of colonies will be distinct. Fortunately, the standard error is still well behaved over a broad space in fraction of regions occupied (Appendix), meaning that plate counts into the “uncrowded” range will still produce good estimates with this method. In fact, this produces a result equivalent to that of the Poisson method in the fully uncrowded regime. However, the MPN method is most useful as plating densities encroach into the crowded regime, allowing precise and accurate estimation of CFU density from plates that would provide severely biased estimates using a naive Poisson model.

### Utility of the Models

Here we demonstrate the relative utility of each model for estimation of CFU density from simulated data. First, we can use the crowding-explicit binomial sampling model described in the previous section, to estimate bias due to crowding, and to demonstrate the importance of choosing an appropriate cutoff *M*, below which plates are considered to be uncrowded and countable. To do so, we solve the crowded binomial model in Eq. 13 for *dV* with respect to the average number of colonies ⟨*n*⟩ and the number of colony-sized regions on a plate *N*. Doing so we find 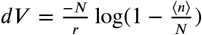. We can substitute this into the Poisson estimator and find:

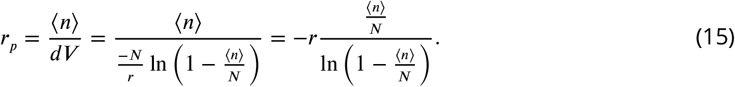

Let us define the ratio of expected colony number to the number of colony-sized regions as 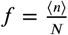. This ratio represents the amount of crowding, where a value of 1 is the maximum crowding and a value close to zero is in the uncrowded regime. Expressing the previous expression in terms of the crowding we see

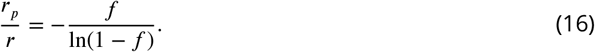

This ratio indicates how close the estimated CFU concentration is to the true concentration. A ratio of 1 tells us that we have an unbiased estimator, whereas a ratio of less than 1 tells us we are underestimating the CFU density. We plot this expression in Fig. 2 to show how the simple Poisson estimator underestimates the actual concentration as a function of crowding, *f*. After a crowding value of *f* = 0.2 the Poisson estimator starts to be significantly biased, undershooting the true value by about 10%. This has implications for the value used in the Poisson model with a cutoff. The cutoff should be chosen such that the bias is not greater than the experimenters targeted precision. For example, if a bias must be less than 10%, then a cutoff of about 20% of the total plate capacity should be used. In the case of a 10 cm plate with an estimated 5000 1mm diameter colony-sized regions, this corresponds to a cutoff of *M* = 1000, whereas the more typical cutoff of *M* = 300 provides an essentially unbiased estimate (bias 3%), but this results in a large statistical fluctuation of 5.8%. In the case of 6mm grid grid on a 10cm by 10cm plate, there are roughly 200 grid regions in a plate. Thus an *M* = 40 would be appropriate to achieve the bias less than 10%, and a threshold of *M* = 12 colonies is required to reduce bias to 3% for colonies of this size. At these thresholds, the statistical error would be 15.8%.

To compare the performance of the different estimators discussed here, we simulated 1000 experiments and applied each of our estimators to the resulting data. Data for each experiment was modeled using the binomial crowding model with *r* = 10^5^, *V* = 0.2, *N* = 5000, and dilution values (0.1, 0.1, 0.01, 0.01, 0.001, 0.001). This corresponds to two replicates for each dilution in a tenfold dilution experiment. An example set of colony counts corresponding to these dilutions is (1705, 1629, 196, 181, 21, 21). The first two dilutions are in the over-crowded regime and the last two dilutions are in the dilute uncrowded regime. The traditional methods (“pick-the-best”, averaging, segment averaging) and Poisson with a cutoff will discard the first two counts as too many to count, while the other methods will use their numeric values. The resulting distributions are plotted in Fig. 3.

**Figure 3.**
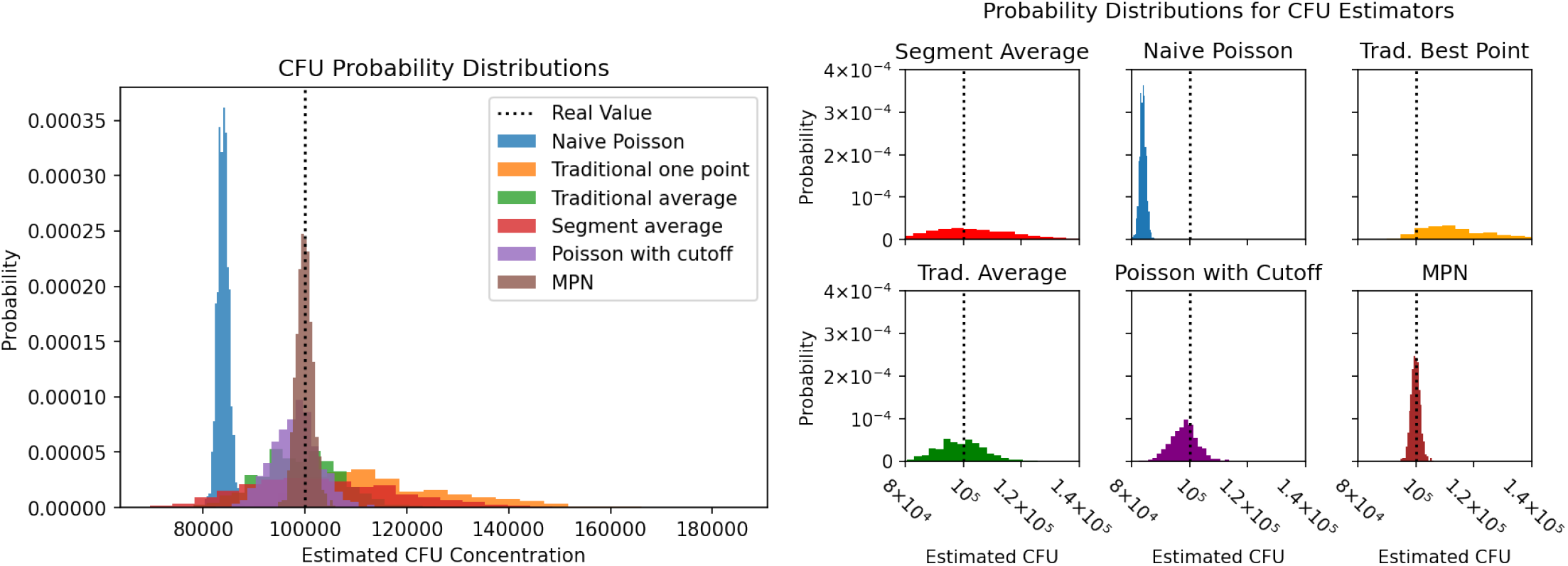
The probability distributions of estimated CFU concentrations from different estimators generated from 1000 independent numerical experiments with dilutions 0.1, 0.01, 0.01, 0.001, 0.001, *r* = 100000, *V* = 0.2, *N* = 5000. Here the Segmented average, naive Poisson, “pick the best”, traditional average, Poisson with cuttoff, and MPN methods are compared. The MPN method demonstrates the best combination of high

The results show that the MPN (most probable number) method is unbiased and has the highest degree of accuracy. The Poisson with a cutoff (which always discards counts from the least-diluted samples in these outputs) is nearly unbiased, whereas the naive Poisson is biased down due to inclusion of “crowded” data. The naive Poisson has a similar variance as that of the MPN because both are using all the data points. However, the measure around which the naive Poisson estimator varies is incorrect due to this bias. With the Poisson estimator, increasing accuracy comes at a cost in precision; the Poisson with cutoff has roughly twice the standard error of the MPN method due to the fact that it does not use all the data and throws out the first two counts of each experiment. Next, the the traditional averaging method (***USDA, 2015***) has roughly five times the standard error of the MPN method, due to the fact that it gives lower-precision measurements the same weight as higher-precision large counts in the uncrowded regime. However, it is unbiased. If there are technical replicates, pick-the-best (choosing the largest number of counts in the countable range, over multiple technical replicates at each dilution) is a biased estimator (overestimating CFUs) and has a standard error roughly ten times that of the MPN method. (Pick-the-best where the best count from *each* technical replicate is used is equivalent to Poisson with a cutoff, with some loss of precision due to discarding of small counts.) Segment averaging (here, counting one-quarter of the plate, and assuming perfect segmenting such that exactly one-quarter of the colonies are counted) resulted in an unbiased estimator with the largest standard error, roughly 13 times the standard error of the MPN method.

These simulations show that the MPN method produces the most precise results and is unbiased. However, the Poisson with a cutoff is a close second, also with high accuracy and precision and with the advantage of being practical to calculate by hand. The bias of the naive Poisson (using all data) serves as a warning: if counts are not in the uncrowded regime, the Poisson assumptions do not apply, and an estimator using only number of colonies counted at each dilution will underestimate the CFU density in the original sample. Other standard estimators (averaging, segment averaging) using the same data required for the Poisson estimator show universally poorer precision than Poisson with a cutoff and cannot be recommended.

## Conclusion

We have presented several methods for estimating CFUs and we have provided a calculator for these estimators available on `Hugging Face` spaces, named `CFUestimator` (***Martini, 2023***). In practice, the choice of method will depend on the precision required for the estimate of CFU density. For experiments with reasonably large expected effect size, the simplest mathematically admissible method - the Poisson estimator with a cutoff - is perfectly valid, as long as the dilutions are chosen appropriately to ensure all measurements are in the countable range. Broadly speaking, addition of unbiased data will improve the precision of an estimator. Historically, technical replicates have been used for this purpose - even technical duplication is sufficient to markedly reduce variance of the estimated CFU density, although triplicate plating is preferred to safeguard against accidents and outliers (***Weenk, 1992***) (also see Appendix). The Poisson model allows data from technical replicates to be combined into a single mathematically interpretable estimator with definable properties - specifically, a maximum likelihood estimator, which should be an unbiased and minimally variable estimator for the true value. This is as opposed to averaging (***USDA, 2015***), which produces an estimate whose properties are not well defined. The Poisson method also allows the investigator to incorporate data from dilutions with too few counts, *in addition to* (not in place of) data from countable wells in the same dilution series - by effectively re-weighting the contribution of these wells by the total volume of original suspension that they contain, these data can be used to improve the accuracy of the estimator even though their sampling variance is high.

The correspondence shown here between using tubes and gridding a plate into subsections based on colony area allows the usage of estimator techniques typically used for quantal-based measurements of CFU density, specifically the MPN, where positive growth events (e. g., colonies) are explicitly considered to represent *one or more* originating cells. These techniques have a long history in environmental surveillance microbiology, and statistically well-founded techniques are readily available for analysis of such data (***Cochran, 1950; Garthright, 1993; Loyer and Hamilton, 1984***). If an experimentalist wants tighter bounds for an estimated CFU count, the MPN provides a very low-variance, unbiased estimator at the cost of some extra steps. This estimator allows the experimentalist to incorporate data from normally uncountable (TMTC) plates as well as counts from uncrowded plates, maximizing the amount of information that can be gleaned from a dilution series.

The MPN model requires an estimate of the maximum number of colonies that can be packed into the growth area for each sample; we show (Appendix) that it is better to over-estimate this maximum than to under-estimate it. If the patch size on a plate is correctly chosen to be around the size of a typical colony, even a spot-plating assay on a 10 by 10 cm plate is equivalent to running hundreds of tubes in parallel. Further, it is necessary to estimate the number of occupied regions in the growth area. In or near the uncrowded regime, this will be equivalent to the number of counts. However, this method does not require that all colonies are individually countable - instead, image analysis (***Chiang et al*., *2015; Brugger et al*., *2012; Bewes et al*., *2008***) can be used to estimate both the size of an individual colony and the fraction of total area occupied by colony growth. The MPN estimator can therefore potentially provide accurate, precise estimates of CFU density for plates where exact counts cannot be obtained. However, colony size varies across different microorganisms as well as across culture conditions (media type, agar percentage, pad thickness, plate drying time and conditions, growth temperature and atmosphere, etc.) and incubation time on plates, meaning that the size range of colonies may be different even across plates within a single experiment (***Savage and Halvorson, 1941; Chacón et al., 2018***). This added complication of properly choosing a grid size or determining the typical size of a colony means that application of the MPN will most likely require parameters estimated for the specific experiment being analyzed. Further, the fact that colony size can decrease under crowding means that heavily-crowded plates or plate regions, where few or no distinct colonies are visible, may have very different “average” colony sizes than the same microbes in a less-crowded area. While theory suggests that the MPN estimator will be most precise when the majority of colony-sized locations are occupied (***Strijbosch et al*., *1987***), also see Appendix, this practical limitation suggests that use of the MPN on plate count data will become less accurate with extremes of crowding, and that the best use of the MPN is likely to be in the liminal region between the technically uncrowded and the physically uncountable, where most to all growth is in the form of distinct, countable colonies but crowding produces a measurable bias in these counts.

## Acknowledgements

This work was funded, in part, by NSF Grant No. 2014173. Ilya Nemenman was additionally supported by the Simons Investigator program.

## Appendix 1

### Introduction

In this Appendix, we mathematically derive the Maximum Likelihood Estimators for CFU concentration presented in the Main Text. Section Poisson Model reviews the basic Poisson model for CFU estimation, Section Poisson Model with a Cutoff derives the Poisson model with a cutoff, and Section Binomial Model of Crowding (MPN) derives the MPN method applied to crowded plate data. Table 1 defines a dictionary of variable names and symbols, and Table 2 presents a summary of the estimators discussed in the Main Text. Finally, Section Bias of the Poisson estimator applied to crowded data, discusses the bias of the naive Poisson estimator for crowded data.

We briefly summarize the problem and the relevant variables as follows: In a typical experiment, there is a stock liquid with an unknown concentration of microbes *r*. This initial liquid is divided between multiple plates or tubes and can be diluted by different factors. In the case of plates, this is with the goal of producing low enough concentrations, such that any resulting colonies can then easily be counted. Similarly, in tubes, the aim is to have multiple tubes, such that some contain viable stock, and some do not. Each plate and/or tube contains the same volume of liquid *V*. However, the concentration has been diluted by a factor *d*_*k*_ = *V*_*k*_/*V*, where *V*_*k*_ is the amount of liquid from the original sample used in the dilution for the *k*th tube or plate. This implies that, for an experiment *k*, we have a volume *V*_*k*_ of liquid with density *d*_*k*_*r* cells per unit volume. Colonies are allowed to grow on each plate, and the total number of colonies is counted as *n*_*k*_.

All the following methods assume that the microbes are randomly distributed throughout each sample, that microbes do not cluster or attract one another, that their growth is independent of one another, and that the medium is not selective.

### Poisson Model

The Poisson model of population counts of microbes assumes that there is a uniform population density *r* of microbes per unit volume in an initial volume *V*_0_ of liquid. The liquid is well mixed and will result in *n*_*k*_ colonies, where *n*_*k*_ is Poisson distributed with a parameter *λ* = *rd*_*k*_*V*. That is, the average number of colonies per experiment is *rd*_*k*_*V* with variance *rd*_*k*_*V*. Each experimental volume is spread uniformly across entire plates resulting in cells being randomly distributed across the plate. The locations of the initial cells are uniformly random and are independent of the locations of where other cells landed. Additionally, it is assumed that each microbe will grow up into a full colony, which then can be counted independently of all other colonies. The Poisson model assumes no crowding.

The likelihood that an experiment *k*, with dilution rate *d*_*k*_, has *n*_*k*_ colonies is

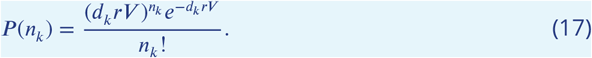

The combined likelihood of all experiments is

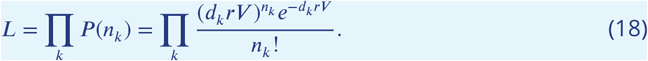

The value of concentration, *r*, that maximizes this likelihood or equivalently the value of *r* that maximizes the log likelihood, is known as the maximum likelihood estimator (*r*_mle_). The maximum likelihood estimator of concentration is an unbiased estimator given the data follows the same probability distribution as given by the model. In reality, the data follows a different distribution.

To find *r*_mle_, we find the maximum of the log-likelihood by setting its derivative w. r. t. *r* to zero:

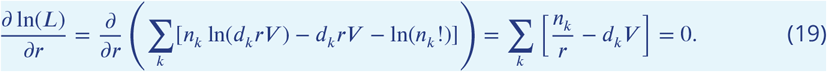

This implies

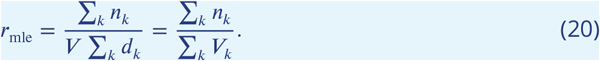

The standard error of the maximum likelihood estimator can be calculated from the second order derivative of the log-likelihood w. r. t. *r*:

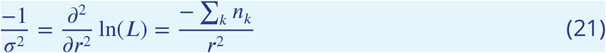

After some algebra, the variance of the mle becomes

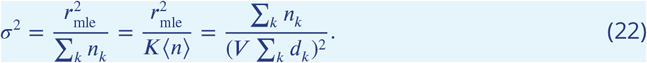

### Poisson Model with a Cutoff

In the Poisson model with a cutoff, it is assumed that colonies are still distributed via a Poisson distribution. However, above a critical colony count, *M*, the colonies start to merge and the experimenter will not count the colonies, instead assigning the category of “too crowded to count”.

The probability that colony counts are above the threshold *M* is calculated as

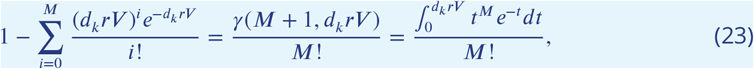

where *γ*(*N* + 1, *d*_*k*_*rV*) is the lower incomplete gamma function. We use indicator functions to write the likelihood of colony counts, where the indicator function *I*(*n* < *M*) is 1 when *n* < *M*, and 0 otherwise. Similarly, *I*(*n* > *M*) is 1 when *n* > *M*, and 0 otherwise. For the purpose of writing the likelihood, if the number of colonies *n*_*k*_ is greater than the threshold *M*, we will assign it the count of *M* + 1 which will act as a special category corresponding to all plates that are too crowded to count. The likelihood is then:

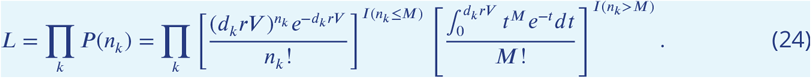

We find the value of *r* that maximizes the the log likelihood in a manner similar to the previous section:

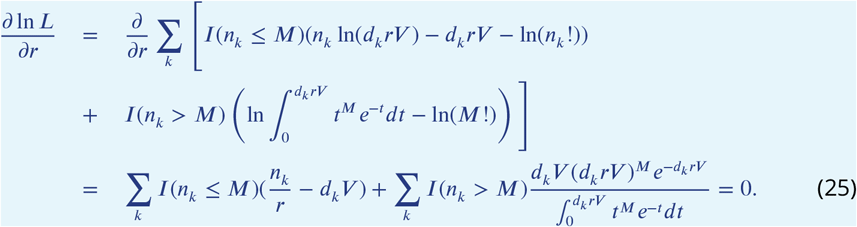

This expression can be solved for numerically for *r*.

There are several limits of this equation that can be easily understood. First, in the un-crowded dilution regime, where *d*_*k*_*rV* ≪ 1, the MLE estimator of *r* becomes

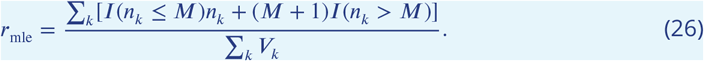

A more useful limit is when there are no data points where *d*_*k*_*rV* ≈ *M* and the data falls into two categories, either *d*_*k*_*rV* ≪ *M* or *d*_*k*_*rV* ≫ *M*. In this limit, the second term in equation 25 corresponding to colony counts larger than *M* can be ignored. This is equivalent to the normal Poisson solution, but with all colony counts larger than the maximum *M* being ignored. The resulting solution is:

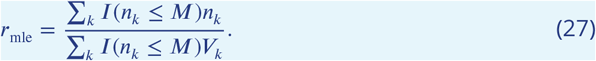

In general, this simple approximate formula gives a result almost identical to the exact numerical solution for *r* in this regime.

### Binomial Model of Crowding (MPN)

To account for crowding,we will divide the plate into *N* regions each the size of a full colony. We then make the assumption that if more than one bacterium lands in one of these regions, the colonies that would form from these cells will grow together and be counted as one colony. For each region, the number of cells landing in that region will be Poisson distributed with parameter 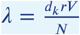. The probability of 0 cells landing in a region is 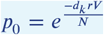, and the probability of more than one cell landing in a region is 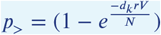.

The number of colonies observed will be binomially distributed

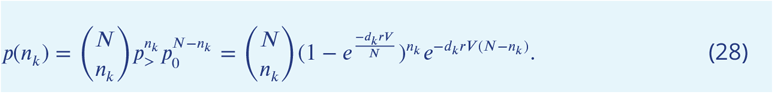

Again, we can find the *r* that maximizes the log-likelihood for this crowding model.

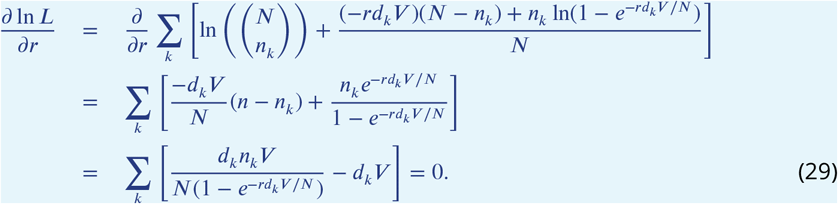

We can numerically solve the last equation to find *r*. One limit that is exactly solvable is when *d*_*k*_ = *d*. If we had *K* trials where all the dilutions where exactly the same, the best estimate for *r* becomes 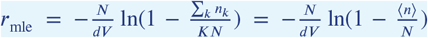. This is actually directly related to the expected number of colonies found from the binomial distribution, which is ⟨*n*⟩ = ⟨*n*⟩ *N* (1 − *e*^−*rdV* /*N*^). Similarly, the binomial distribution will have a variance of the expected number of colonies of *N*(1 − *e*^−*rdV*/ *N*^)(*e*^−*rdV*/ *N*^).

In the small concentration limit *rdV* /*N* ≪ 1, the mean and the variance are approximately the same, *rdV*, which would be the case for the Poisson distribution. This shows that the crowding model reduces to the Poisson model for low concentrations and low colony counts. However, if you keep to the next highest order in our small parameter, we see that the mean and variance are actually different from one another, namely 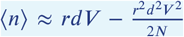 and 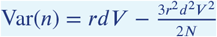.

Previous work (***Ben-David and Davidson, 2014***) has modeled crowding using the shifted Poisson distributions. They claimed the variance would be the same as if there was no crowding, and the mean would be shifted down due to colonies merging together. However, this is inconsistent with our model of crowding. In fact, both the mean and the variance are shifted relative to what their true values would have been if the distributions were purely Poisson. The reason that the variance of large colony counts is also shifted downwards is that we have an upper bound on the total number of colonies, above which the estimator cannot fluctuate since the colonies would merge and be counted as single colonies. In other words, the use of a shifted Poisson distribution is a fine approximation, but the variance— and not just the mean—must also be adjusted.

We can find the error associated with the maximum likelihood estimator for *r* as before, by finding the second order derivative of the log likelihood with respect to *r*:

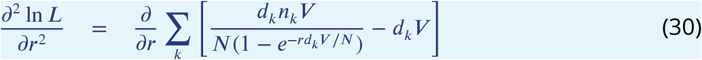

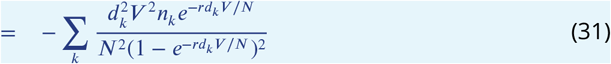

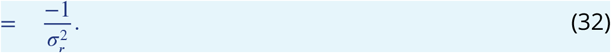

Switching again to the case where we are conducting *K* trials at the same dilution *d*_*k*_ = *d*, we find 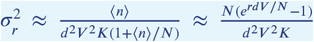. We can attempt now to find if there is an optimal dilution rate that would give us the smallest uncertainty. For this, let 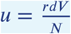. Then the variance becomes 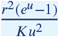. The function 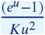 has a minimum when *u* ≈ 1.594, which implies the optimal dilution *d*_opt_ = 1.594*N*/(*rV*). Alternatively, the error is minimized when the average number of colonies is roughly 80% of the maximum total number of colonies possible.

We estimate the size of a region for applying the MPN estimator to plates as the typical size of a colony, so that the maximum number of regions in a plate, *N*, is the ratio of the plate area to the typical colony size area. To understand how sensitive MPN is to this choice, we simulated data from the binomial crowding model with fixed *N*_true_ = 5000 and applied the MPN estimator to the simulated data, while varying the maximum estimated number of colonies *N*. We generated data for *r* = 100000, *V* = 0.2, *N* = 5000, and dilution values *d*_*k*_=0.1, 0.1, 0.01, 0.01, 0.001, 0.001. Figure shows the functional dependence of the estimated value on the guess for *N*. This plot illustrates that the estimator reduces to the naive-Poisson estimator as the max number of colonies approaches infinity. It also shows that, if the measured number of colonies from an experiment is close to the guessed *N*, the estimator diverges. Indeed, there are many concentrations that correspond to a fully crowded plate. This analysis shows that it is better to use an overestimate for the max number of colonies than an underestimate. An overestimate will, at worst, result in the estimate given by the Naive Poisson estimator. An underestimate, on the other hand, will result in greatly overestimating the role that crowding plays and will bias the estimate of the concentration upwards.

**Appendix 1—figure 1.**
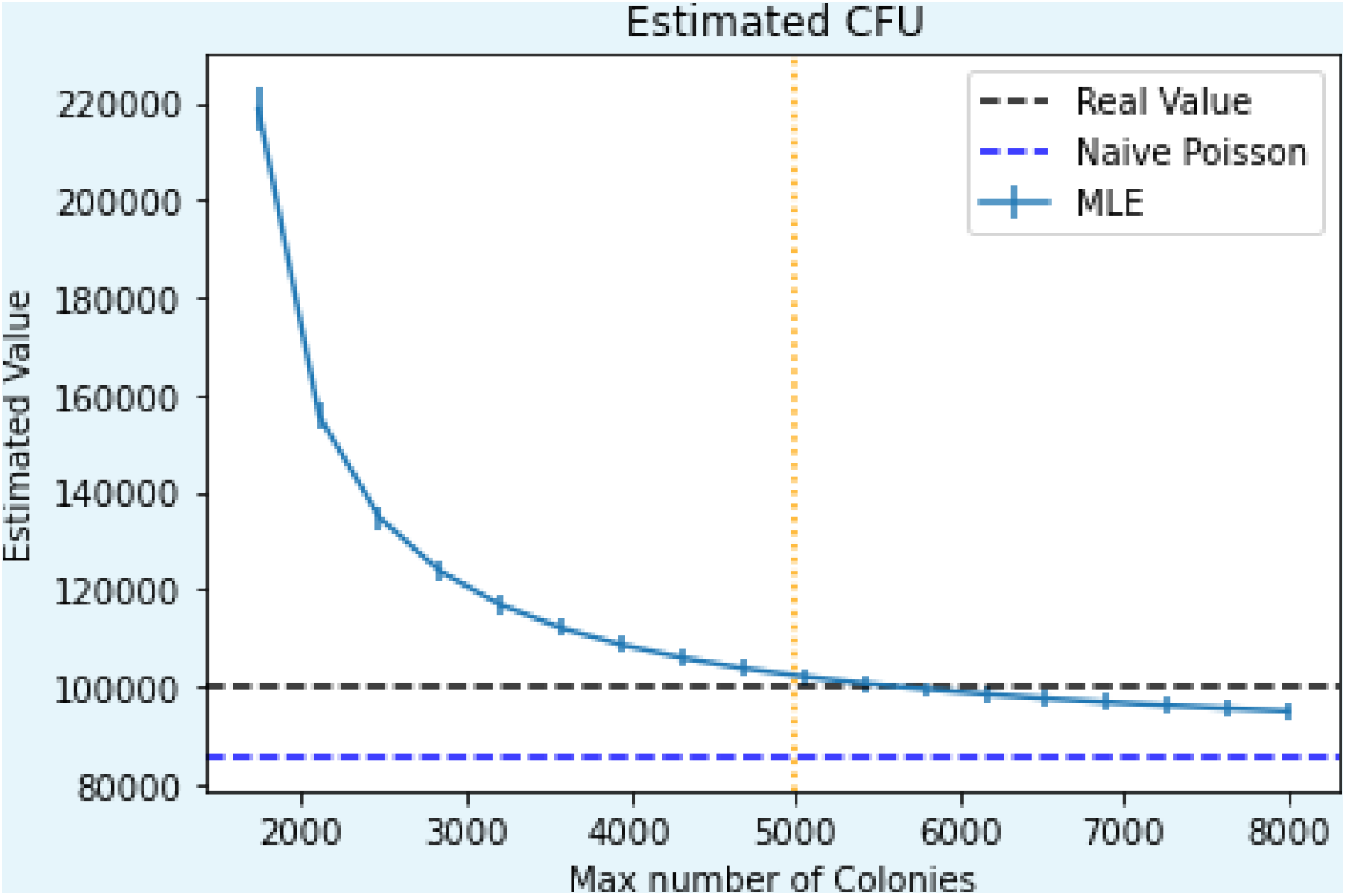
The MPN estimator as a function of an estimated maximum number of colonies, *N*, when the true maximum number is 5000.

### Summary of Methods

Table 1 summarizes variable names, their corresponding symbols, and any relevant notes about the variables. Table 2 summarizes how to calculate different estimators and the corresponding standard error for each estimator.

**Appendix 1—table 1.** A table summarizing variable names and meanings

| Variable Name | Symbol | Notes |
| --- | --- | --- |
| CFU concentration | $r$ | The concentration of microbes (counts per volume) |
| volume of sample | $V$ | The total volume of a sample |
| original sample volume | $V_k$ | The volume of the original sample used for sample $k$ |
| number of counts | $n_k$ | The number of colony counts for sample $k$ |
| dilution factor | $d_k$ | The dilution factor $\frac{V_k}{V_0}$ sample $k$ |
| cutoff threshold | $M$ | The threshold of colonies, above which there are crowding effects |
| maximum number of colonies | $N$ | The maximum number of colony-sized regions in a plate, $N \approx \frac{A_{\text{plate}}}{A_{\text{colony}}}$ |

**Appendix 1—table 2.** A table summarizing the estimator methods along with their corresponding standard errors.

| Estimator Name | Estimator | Standard Error |
| --- | --- | --- |
| Naive Poisson | $\frac{\sum_k n_k}{\sum_k V_k}$ | $\sqrt{\frac{r_{\text{mle}}^2}{\sum_k n_k}}$ |
| Poisson with cutoff | $\frac{\sum_k I(n_k \leq M) n_k}{\sum_k I(n_k \leq M) V_k}$ | $\sqrt{\frac{r_{\text{mle}}^2}{\sum_k I(n_k \leq M) n_k}}$ |
| Sophisticated Poisson with cutoff | $\sum_k I(n_k \leq M) \left( \frac{n_k}{r} - d_k V \right) +$ $\sum_k I(n_k > M) \frac{d_k V (d_k r V)^M e^{-d_k r V}}{\int_0^{d_k r V} t^M e^{-t} dt} = 0$ | $\sqrt{\frac{r_{\text{mle}}^2}{\sum_k I(n_k \leq M) n_k}}$ |
| MPN | $\sum_k \frac{d_k n_k}{N(1 - e^{-rd_k V/N})} = \sum_k d_k$ | $\sqrt{\frac{\langle n \rangle}{d^2 V^2 K(1 + \langle n \rangle/N)}}$ |

### Bias of the Poisson estimator applied to crowded data

We use the above binomial model of crowding to get an estimate of the bias of the simple Poisson model applied to crowded data collected at the same dilution *d*_*k*_ = *d*. In this situation, the Poisson estimator is 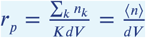. We substitute into this expression the expected number of colonies from the binomial crowding model to find how well the Poisson estimator works under these conditions.

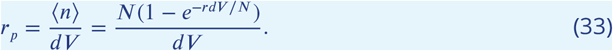

Similarly we find how the standard error should behave as a function of dilution.

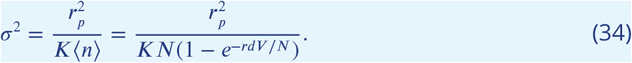

We simulated data from the binomial crowding model and applied the Poisson estimator to the data generated from the model. We generated data for *r* = 100000, *V* = 0.2, *N* = 5000, and *d* ranging from 10^−4^ to 10^−1^. The following Figs. 2, 3, 4 show the theoretical curves along with the predictions for specific data generated at the given dilutions and different number of replicate measurements. Notice how very low dilutions (corresponding to a few counts per plate) give unbiased but higher-variance estimates. Also note that, up to a dilution of about 5 · 10^−2^, both estimators give relatively unbiased results. However, for dilution factors approaching 5·10^−2^ (corresponding to about 1000 colonies per plate), the Poisson estimator starts to give very biased estimates undershooting the true value of the concentration.

**Appendix 1—figure 2.**
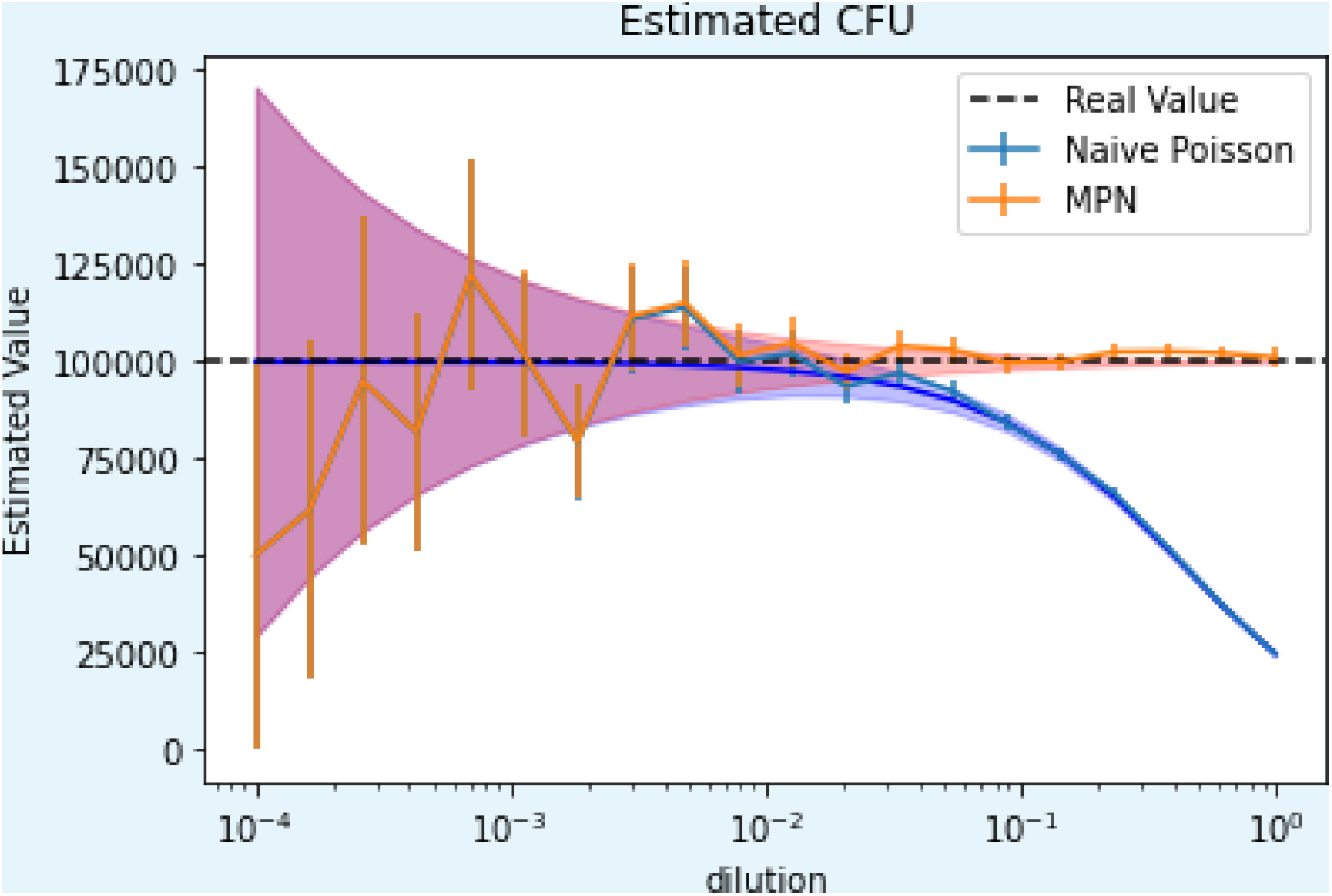
The naive Poisson estimator and the MPN estimator for one realization per dilution of data drawn from a binomial crowding model. Shaded regions correspond to the theoretical standard error of both estimators.

**Appendix 1—figure 3.**
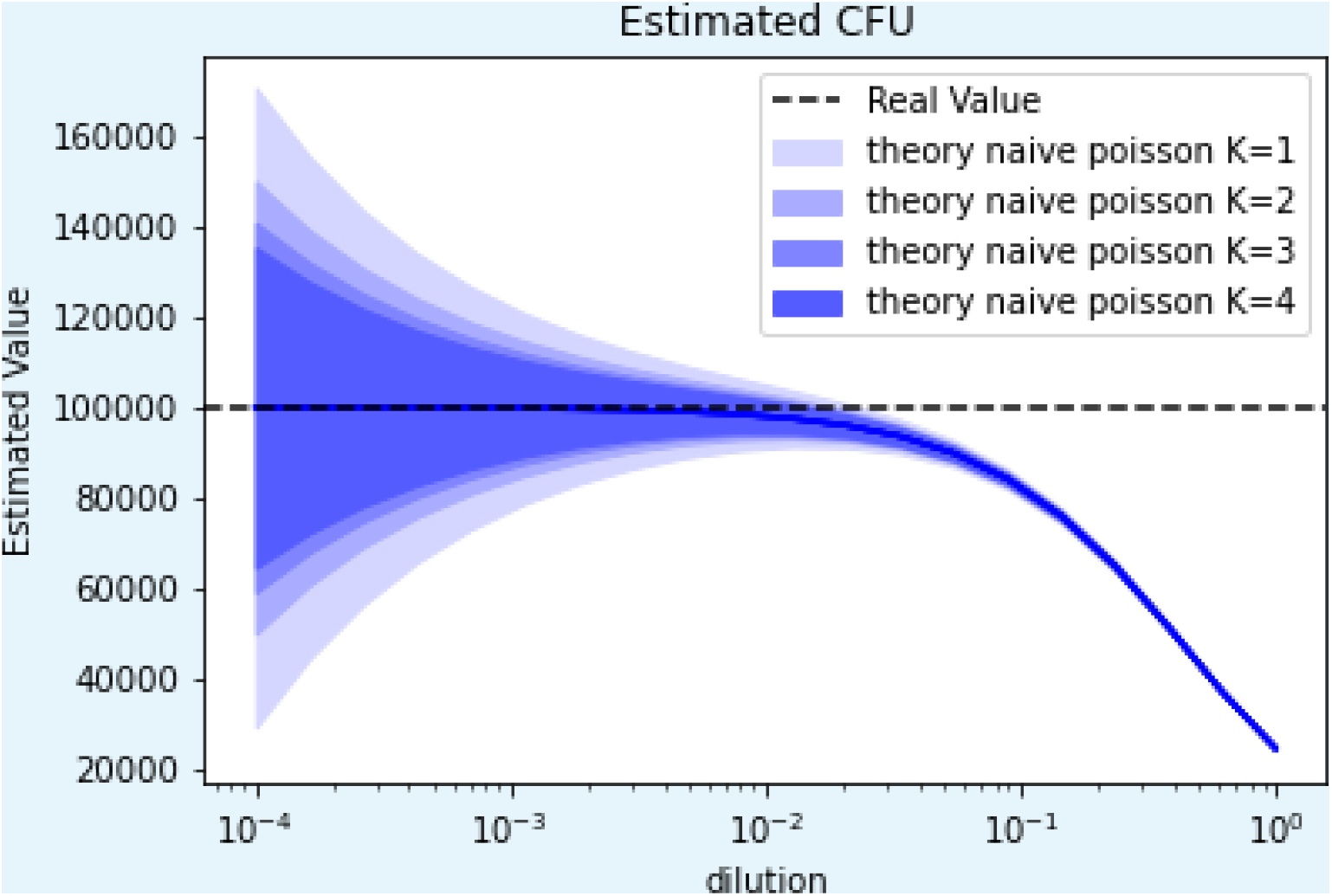
The naive Poisson estimator with standard error for replicates *K* = 1, *K* = 2, *K* = 3, and *K* = 4.

**Appendix 1—figure 4.**
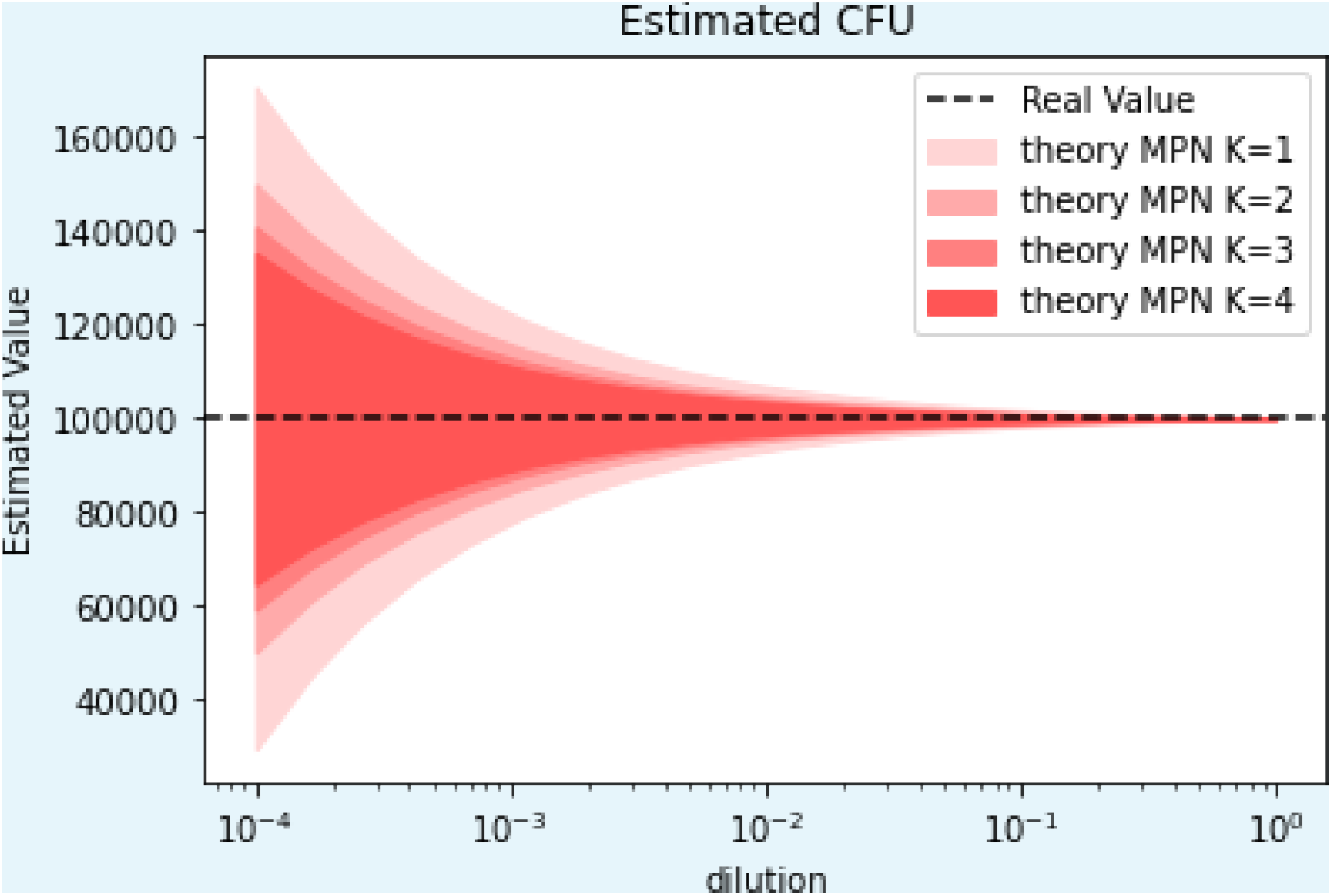
The MPN estimator with standard error for replicates *K* = 1, *K* = 2, *K* = 3, and *K* = 4.

Another way to view this data is to solve the crowded binomial model for *dV* with respect to the average number of colonies and maximum number of colonies allowed. Doing so we find 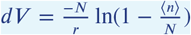. We can substitute this into the Poisson estimator and find:

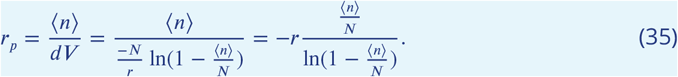

Let us define the ratio of the expected colony number to the maximum colony number as 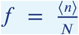. This ratio represents the amount of crowding, a value of 1 is the maximum crowding and a value close to zero is in the uncrowded regime. Expressing the previous expression in terms of the crowding, we obtain:

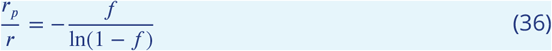

This ratio indicates how close the estimated concentration is to the true concentration. A ratio of 1 tells us that we have an unbiased estimator, a ratio of less than 1 tells that we are underestimating the true value of the concentration. We plot this expression in Fig. 2 in the Main Text to show how the Poisson estimator underestimates the actual concentration as a function of crowding, *f*. After a crowding value of *f* = 0.2 the naive Poisson estimator starts to be significantly biased, undershooting the true value by about 10%. This has implications for the value used in the Poisson model with a cutoff. The cutoff should be chosen such that the bias is not greater than the experimenters targeted precision. For an error less than 10%, a cutoff of about 20% of the maximum number of colonies should be used. In the case of a maximum of 5000 colonies, this corresponds to a cutoff of *M* = 1000.

## Notes

### Competing Interest Statement

The authors have declared no competing interest.

### Summary of Updates

Article has been reformatted and now includes the supplementary information as an appendix.

## References

Alexander M. Most probable number method for microbial populations. Methods of Soil Analysis: Part 2 Chemical and Microbiological Properties. 1983; 9:815–820.

Ben-David A, Davidson CE. Estimation method for serial dilution experiments. Journal of microbiological methods. 2014; 107:214–221.

Bewes J, Suchowerska N, McKenzie D. Automated cell colony counting and analysis using the circular Hough image transform algorithm (CHiTA). Physics in Medicine & Biology. 2008; 53(21):5991.

Blodgett R, BAM appendix 2: Most probable number from serial dilutions. FDA; 2020. https://www.fda.gov/food/laboratory-methods-food/bam-appendix-2-most-probable-number-serial-dilutions.

Breed RS, Dotterrer W. The number of colonies allowable on satisfactory agar plates. Journal of bacteriology. 1916; 1(3):321–331.

Brewster JD. A simple micro-growth assay for enumerating bacteria. Journal of Microbiological Methods. 2003; 53(1):77–86.

Brugger SD, Baumberger C, Jost M, Jenni W, Brugger U, Mühlemann K. Automated counting of bacterial colony forming units on agar plates. PloS one. 2012; 7(3):e33695.

Canales RA, Wilson AM, Pearce-Walker JI, Verhougstraete MP, Reynolds KA. Methods for Handling Left-Censored Data in Quantitative Microbial Risk Assessment. Applied and Environmental Microbiology. 2018 Oct; 84(20):e01203–18. https://journals.asm.org/doi/full/10.1128/AEM.01203-18, doi: 10.1128/AEM.01203-18, publisher:American Society for Microbiology.

Chacón JM, Möbius W, Harcombe WR. The spatial and metabolic basis of colony size variation. The ISME Journal. 2018 Mar; 12(3):669–680. https://www.nature.com/articles/s41396-017-0038-0, doi: 10.1038/s41396-017-0038-0.

Chiang PJ, Tseng MJ, He ZS, Li CH. Automated counting of bacterial colonies by image analysis. Journal of microbiological methods. 2015; 108:74–82.

Christen JA, Parker AE. Systematic statistical analysis of microbial data from dilution series. Journal of Agricultural, Biological and Environmental Statistics. 2020; 25(3):339–364.

Clarke K, Owens N. A simple and versatile micro-computer program for the determination of ‘most probable number’. Journal of Microbiological Methods. 1983; 1(3):133–137.

Cochran WG. Estimation of bacterial densities by means of the” most probable number”. Biometrics. 1950; 6(2):105–116.

Epa C. Evaluation of Options for Interpreting Environmental Microbiology Field Data Results having Low Spore Counts. Environmental Protection Agency; 2014.

Fisher R, Thornton H, Mackenzie W. THE ACCURACY OF THE PLATING METHOD OF ESTIMATING THE DENSITY OF BACTERIAL POPULATIONS: WITH PARTICULAR REFERENCE TO THE USE OF THORNTON’S AGAR MEDIUM WITH SOIL SAMPLES. Annals of Applied Biology. 1922; 9(3-4):325–359.

Garthright WE. Bias in the logarithm of microbial density estimates from serial dilutions. Biometrical Journal. 1993; 35(3):299–314.

Gijbels I. Censored data. WIREs Computational Statistics. 2010; 2(2):178–188. https://onlinelibrary.wiley.com/doi/abs/10.1002/wics.80, doi: 10.1002/wics.80, _eprint: https://onlinelibrary.wiley.com/doi/pdf/10.1002/wics.80.

Gronewold AD, Wolpert RL. Modeling the relationship between most probable number (MPN) and colonyforming unit (CFU) estimates of fecal coliform concentration. Water research. 2008; 42(13):3327–3334.

Haas CN, Rose JB, Gerba CP. Exposure Assessment. In: Quantitative Microbial Risk Assessment John Wiley & Sons, Ltd; 2014.p. 159–234. https://onlinelibrary.wiley.com/doi/abs/10.1002/9781118910030.ch6, doi: 10.1002/9781118910030.ch6,section:6 _eprint: https://onlinelibrary.wiley.com/doi/pdf/10.1002/9781118910030.ch6.

Halvorson H, Ziegler N. Application of statistics to problems in bacteriology: I. A means of determining bacterial population by the dilution method. Journal of Bacteriology. 1933; 25(2):101–121.

Harris R, Sommers L. Plate-dilution frequency technique for assay of microbial ecology. Applied microbiology. 1968; 16(2):330–334.

Hedges AJ. Estimating the precision of serial dilutions and viable bacterial counts. International journal of food microbiology. 2002; 76(3):207–214.

Herigstad B, Hamilton M, Heersink J. How to optimize the drop plate method for enumerating bacteria. Journal of microbiological methods. 2001; 44(2):121–129.

Horvitz DG, Thompson DJ. A generalization of sampling without replacement from a finite universe. Journal of the American statistical Association. 1952; 47(260):663–685.

Hurley MA, Roscoe M. Automated statistical analysis of microbial enumeration by dilution series. Journal of applied bacteriology. 1983; 55(1):159–164.

Jennison MW, Wadsworth GP. Evaluation of the errors involved in estimating bacterial numbers by the plating method. Journal of Bacteriology. 1940; 39(4):389–397.

Jett BD, Hatter KL, Huycke MM, Gilmore MS. Simplified agar plate method for quantifying viable bacteria. Biotechniques. 1997; 23(4):648–650.

Johnson EA, Brown Jr BW. The Spearman estimator for serial dilution assays. Biometrics. 1961; p. 79–88.

Jones P, Mollison J, Quenouille M. A Technique for the Quantitative Estimation of Soil Micro-organisms: With a Statistical Note by. Microbiology. 1948; 2(1):54–69.

Kirchman D, Sigda J, Kapuscinski R, Mitchell R. Statistical analysis of the direct count method for enumerating bacteria. Applied and Environmental Microbiology. 1982; 44(2):376–382.

Loyer MW, Hamilton MA. Interval estimation of the density of organisms using a serial-dilution experiment. Biometrics. 1984 Dec; 40(4):907–916. doi: 10.2307/2531142,mAG ID: 2317336612.

Luria SE, Delbrück M. Mutations of bacteria from virus sensitivity to virus resistance. Genetics. 1943; 28(6):491.

Martini KM, Calculate Colony Forming Units (CFUs); 2023. https://huggingface.co/spaces/KMichaelMartini/CFUestimator.

McCrady MH. The numerical interpretation of fermentation-tube results. The Journal of Infectious Diseases. 1915; p. 183–212.

Salama IA, Koch GG, Tolley DH. On the estimation of the most probable number in a serial dilution experiment. Communications in Statistics - Theory and Methods. 1978 Jan; 7(13):1267–1281. https://doi.org/10.1080/03610927808827710, doi: 10.1080/03610927808827710, publisher:Taylor & Francis _eprint: https://doi.org/10.1080/03610927808827710.

Savage GM, Halvorson HO. The Effect of Culture Environment on Results Obtained with the Dilution Method of Determining Bacterial Population. Journal of Bacteriology. 1941 Mar; 41(3):355–362. doi: 10.1128/jb.41.3.355-362.1941,mAG ID: 108945167.

Skinner F, Jones P, Mollison J. A comparison of a direct-and a plate-counting technique for the quantitative estimation of soil micro-organisms. Microbiology. 1952; 6(3-4):261–271.

Strijbosch LW, Buurman WA, Does RJ, Zinken PH, Groenewegen G. Limiting dilution assays: experimental design and statistical analysis. Journal of immunological methods. 1987; 97(1):133–140.

Sutton S. Accuracy of plate counts. Journal of validation technology. 2011; 17(3):42–46.

Taylor J. The estimation of numbers of bacteria by tenfold dilution series. Journal of Applied Bacteriology. 1962; 25(1):54–61.

Tremaine SC, Mills AL. Tests of the critical assumptions of the dilution method for estimating bacterivory by microeucaryotes. Applied and environmental microbiology. 1987; 53(12):2914–2921.

USDA, Quantitative Analysis of Bacteria in Foods as Sanitary Indicators. USDA; 2015. https://www.fsis.usda.gov/sites/default/files/media_file/2021-03/MLG-3.pdf.

Weenk GH. Microbiological assessment of culture media: comparison and statistical evaluation of methods. International Journal of Food Microbiology. 1992 Oct; 17(2):159–181. https://www.sciencedirect.com/science/article/pii/016816059290113H, doi: 10.1016/0168-1605(92)90113-H.

